# Functional and genetic adaptations contributing to *Enterococcus faecalis* persistence in the female urinary tract

**DOI:** 10.1101/2023.05.18.541374

**Authors:** Belle M. Sharon, Amanda P. Arute, Amber Nguyen, Suman Tiwari, Sri Snehita Reddy Bonthu, Neha V. Hulyalkar, Michael L. Neugent, Dennise Palacios Araya, Nicholas A. Dillon, Philippe E. Zimmern, Kelli L. Palmer, Nicole J. De Nisco

## Abstract

*Enterococcus faecalis* is the leading Gram-positive bacterial species implicated in urinary tract infection (UTI). An opportunistic pathogen, *E. faecalis* is a commensal of the human gastrointestinal tract (GIT) and its presence in the GIT is a predisposing factor for UTI. The mechanisms by which *E. faecalis* colonizes and survives in the urinary tract (UT) are poorly understood, especially in uncomplicated or recurrent UTI. The UT is distinct from the GIT and is characterized by a sparse nutrient landscape and unique environmental stressors. In this study, we isolated and sequenced a collection of 37 clinical *E. faecalis* strains from the urine of primarily postmenopausal women. We generated 33 closed genome assemblies and four highly contiguous draft assemblies and conducted a comparative genomics to identify genetic features enriched in urinary *E. faecalis* with respect to *E. faecalis* isolated from the human GIT and blood. Phylogenetic analysis revealed high diversity among urinary strains and a closer relatedness between urine and gut isolates than blood isolates. Plasmid replicon (rep) typing further underscored possible UT-GIT interconnection identifying nine shared rep types between urine and gut *E. faecalis*. Both genotypic and phenotypic analysis of antimicrobial resistance among urinary *E. faecalis* revealed infrequent resistance to front-line UTI antibiotics nitrofurantoin and fluoroquinolones and no vancomycin resistance. Finally, we identified 19 candidate genes enriched among urinary strains that may play a role in adaptation to the UT. These genes are involved in the core processes of sugar transport, cobalamin import, glucose metabolism, and post-transcriptional regulation of gene expression.

**IMPORTANCE:** Urinary tract infection (UTI) is a global health issue that imposes substantial burden on healthcare systems. Women are disproportionately affected by UTI with >60% of women experiencing at least one UTI in their lifetime. UTIs can recur, particularly in postmenopausal women, leading to diminished quality of life and potentially life-threatening complications. Understanding how pathogens colonize and survive in the urinary tract is necessary to identify new therapeutic targets that are urgently needed due to rising rates of antimicrobial resistance. How *Enterococcus faecalis*, a bacterium commonly associated with UTI, adapts to the urinary tract remains understudied. Here, we generated a collection of high-quality closed genome assemblies of clinical urinary *E. faecalis* isolated from the urine of postmenopausal women that we used alongside detailed clinical metadata to perform a robust comparative genomic investigation of genetic factors that may mediate urinary *E. faecalis* adaptation to the female urinary tract.

## INTRODUCTION

*Enterococcus faecalis*, a Gram-positive bacterium, is an increasingly frequent cause of urinary tract infection (UTI), especially in complicated or recurrent cases (1–5). While a strong body of work has elucidated virulence mechanisms associated with complicated enterococcal UTI, namely catheter-associated UTI, little is known about how enterococci cause uncomplicated and recurrent UTI (6–11). *E. faecalis* natively inhabits the human gastrointestinal tract (GIT) but is an opportunistic pathogen (12, 13). Its presence in the GIT is proposed to be a predisposing factor to UTI since periurethral contamination by gut bacteria may be an important route of infection (14–16). This idea is supported by a reported association between gut enterococci abundance and *Enterococcus* UTI (14). In addition, premenopausal rUTI patients were found to have greater frequency of infection with endogenous gut microbes, specifically *E. coli* and *E. faecalis* (17–19). Despite the association of *E. faecalis* with rUTI, little is known about the mechanisms by which *E. faecalis* colonizes the urinary tract in the absence of a catheter and the genetic determinants that enable its persistence.

Enterococci demonstrate great adaptability to thrive in stressful environments (13, 20). Urine, as compared to the gut environment, is a nutrient-limited medium characterized by high osmolarity, limited nitrogen and carbohydrate availability, moderate oxygenation, and low pH (21–23). Urine is also considered to be antimicrobial, composed of high concentrations of inhibitory urea and other antimicrobial proteins (23). Survival in the urinary tract despite environmental pressures and antibiotic intervention, suggests *E. faecalis* urine isolates may be specialized genetically or phenotypically. However, understanding of genetic factors necessary for *E. faecalis* growth in the urinary environment is limited.

To date, studies of *E. faecalis* urinary fitness are limited and focus on commensal or blood isolates which are possibly genetically distinct from urinary isolates (24). Previous work to elucidate urinary fitness of *E. faecalis* was conducted on well-studied *E. faecalis* strains OG1RF, V583, and MMH594 grown in pooled urine (25–28). These identified differential expression of genes encoding a sucrose PTS-system, the *lutABC* operon for L-lactate metabolism, an amino acid ABC transporter, and *efaCBA* for manganese scavenging during growth in urine (28). However, no comprehensive study of a collection of clinical urinary *E. faecalis* isolates has been published. Although various virulence factors have been proposed as pathogenicity markers, a virulence genotype to distinguish uropathogenic *E. faecalis* from other *E. faecalis* strains has yet to be proposed (12). Thus, there is a need to conduct a genomic study of urinary *E. faecalis* isolates to identify genetic markers associated with urinary tract colonization.

To gain a deeper understanding of *E. faecalis* adaptation to the urinary environment, we generated a collection of high-quality genomic sequences of clinical urinary *E. faecalis* isolates and compared them to isolates from the human gut and blood – distinct anatomical sites that pose unique evolutionary pressures on *E. faecalis*. Our findings show that urinary strains are diverse, possessing a wide range of plasmid replicon types and exhibiting particularly low antimicrobial resistance rates as determined both by genotypic predictions and phenotypic assessments. We find that vancomycin resistance, which is commonly studied in this genus, was absent from the urine group and intermediate fluoroquinolone resistance phenotypes were common but did not correlate with any known chromosomal mutations or resistance genes. Finally, we identified genes involved in carbohydrate transport and metabolism as well as vitamin B12 import, and post-transcriptional regulation of gene expression as enriched among urinary isolates. Together, this works provides a resource of high quality and well-curated urinary *E. faecalis* genomes and identifies candidate pathways that may be important for *E. faecalis* adaptation to the urinary tract.

## RESULTS

### Collection of urinary E. faecalis strains isolated from the urine of postmenopausal women

While the importance of *E. faecalis* in complicated UTIs like CAUTI has been investigated, the role of *E. faecalis* in uncomplicated UTI or in the microbiome of asymptomatic women is not understood. Nevertheless, *E. faecalis* is commonly isolated from the female urogenital tract (29–31). Indeed, secondary analysis of our recently published metagenomic study of the urinary microbiome of postmenopausal women revealed that *E. faecalis* was present in 57.3% of samples (32). In patients with active rUTI at the time of urine collection, *E. faecalis* was detected in 46.2% of samples. In patients not experiencing infection during urine collection but with history of rUTI, *E. faecalis* was detected in 62.5% of urinary microbiomes (32). These observations strengthen previous reports of the prevalence *of E.* faecalis in the female urogenital tract and highlight the need to understand how *E. faecalis* adapts to this unique environment.

To begin to understand the potential role of genetic adaptation in *E. faecalis* colonization of the female urogenital tract, we sequenced 37 unique *E. faecalis* strains isolated from the urine of postmenopausal women using both Illumina and Nanopore sequencing for the generation of complete or near-complete high quality genome assemblies (**Table S1**). *E. faecalis* strains were isolated from the urine of consenting postmenopausal women that were recruited from a tertiary care center in Dallas, Texas USA between October 2017 and April 2019 (**Figure 1A**, **Table S2**). The 37 women were stratified into four cohorts based on their history of RUTI, UTI symptoms, and urinalysis (UA): Never UTI (no clinical history of symptomatic UTI, n=4), Sporadic (clinical history of UTI, current UTI symptoms, and +UA n=3), Remission (history of RUTI, no current UTI symptoms, and −UA, n=17), and Relapse (history of RUTI, current UTI symptoms, and +UA, n=13). Median patient age was 72 with a median BMI of 27.5. Additional clinically relevant metadata were collected including diabetes status, smoking, electrofulguration history, and prescribed estrogen hormone therapy (EHT), NSAIDs, and antibiotics. In total, 8 women (21.6%) were diabetic, 19 had undergone electrofulguration of trigonitis (51.3%), 18 were prescribed EHT (48.6%), 11 were prescribed NSAIDs (29.7%), and 15 were prescribed antibiotics (40.5%) with 9 of those on current treatment courses. The most prescribed antibiotics were nitrofurantoin followed by TMP-SMX, fluoroquinolone, cephalosporin, and penicillin (**Table S2**). Twenty-five specimens were polymicrobial (67.6%), 7 were dominated by *E.* coli and *E.* faecalis (18.9%), 4 were dominated by *E. faecalis* (10.8%), and one was an *E. faecalis* mono-infection (2.7%). (**Table S2**).

**Figure 1.**
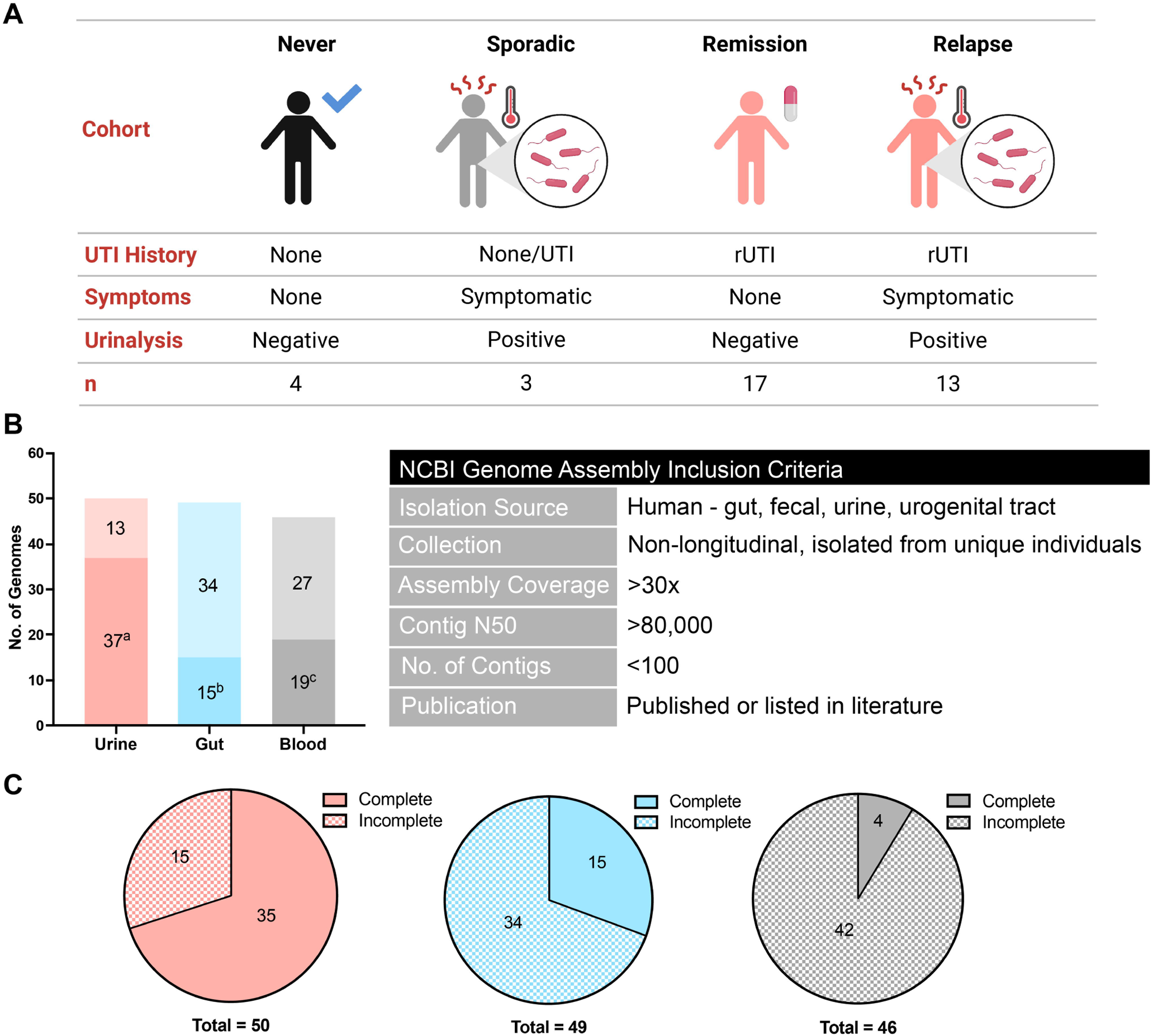
Clinical cohorts and isolated collection of *Enterococcus faecalis*. **(A)** Patient cohorts were stratified based on patient UTI history, symptoms, and urinalysis results at time of specimen collection. n denotes number of isolates. **(B)** Number of isolate genomes from each isolation group. ^a^Isolates sequenced as part of this study. ^b^Isolates from Palacios Araya et al study (72, 73). ^c^Isolates from Van Tyne et al study (24). Genome assembly criteria used for selection of comparator genomes. **(C)** Counts of complete and incomplete genome assemblies in the urine (pink), gut (blue), and blood (grey) isolation groups.

Among the 37 urinary genomes, 33 were complete while 4 remained draft with the most fragmented having <21 highly contiguous fragments (**Table S1)**. As of March 2023, a total of 131 complete genome assemblies of *E. faecalis* isolated from the human host were publicly available on NCBI. Of the 131 complete genomes, only 9 were clearly identified as associated with human urine. This collection, therefore, presents a focused group of high-quality urinary *E. faecalis* genomes. Coupled with clinical metadata, this collection of urinary *E. faecalis* isolates and their complete genomes provides a valuable resource for the field of *E. faecalis* and *E. faecalis* UTI.

### Complete urinary genomes allow comparative analysis of urinary E. faecalis to gut and blood isolates

We hypothesized that urinary *E. faecalis* possesses genetic and phenotypic adaptations that enable its survival in the urinary tract. To test this hypothesis, we curated comparator genomes of diverse *E. faecalis* strains isolated from the urinary tract (n=50), gastrointestinal tract (n=49), and blood (n=46) anatomical niches, which we term “isolation groups” (**Figure 1B, Table S3**). Each isolation group is comprised of publicly available genome assemblies obtained from NCBI that met inclusion criteria described in Materials & Methods (**Figure 1C)**. The majority of urinary strains (74%) were sequenced as part of this study (**Figure 1B**), while 31% of gut isolates were obtained from Dallas, Texas, fecal surveillance (33, 34) and 41% of blood isolates were obtained from a Wisconsin hospital outbreak in the 1980s (24). Proportions of complete assemblies available in the gut (31%) and blood (9%) isolation groups are substantially lower than in the urine group (70%) due to the genome availability at the time of curation (**Figure 1C**).

### Phylogenetic and pangenome analyses reveal high diversity among urinary E. faecalis strains

Comparative analysis of all *E. faecalis* genomes included in this study revealed that urinary strains are diverse (**Figure 2A**). Urinary strains did not form unique clusters in a core gene phylogenetic tree. A large cluster of highly closely related ST6 blood isolates and a large mixed gut and urine cluster of ST179 and ST16 isolates were observed (**Figure 2A**). Small urine-gut clusters as well as blood-gut clusters appear throughout the tree, suggesting closer relatedness between the urinary and gut strains than urinary and blood. The phylogeny largely follows the crude evolutionary estimates provided by Multi-locus Sequence Typing (MLST) (**Figure 2A**). Urinary isolates are characterized by 28 different STs (**Figure 2B**), with ST179 being the most common (14%). Among gut isolates, 27 different STs are present with ST16 being the most common (20%). Interestingly, ST179 and ST16 are single locus variants of each other, differing by a synonymous variation in *xpt*. Blood isolates represent 17 STs and have a strong lineage bias for ST6, with 22 of the 46 isolates belonging to ST6, a hospital outbreak lineage (24). Despite this bias, this collection is representative of the genomic data available for bloodstream *E. faecalis*. We observed that urine isolates had the most unique STs (18) not found among isolates of the other groups, followed by gut (14), and blood (9) and that urine and blood isolates do not share any STs that are absent from the gut group (**Figure 2B**).

**Figure 2.**
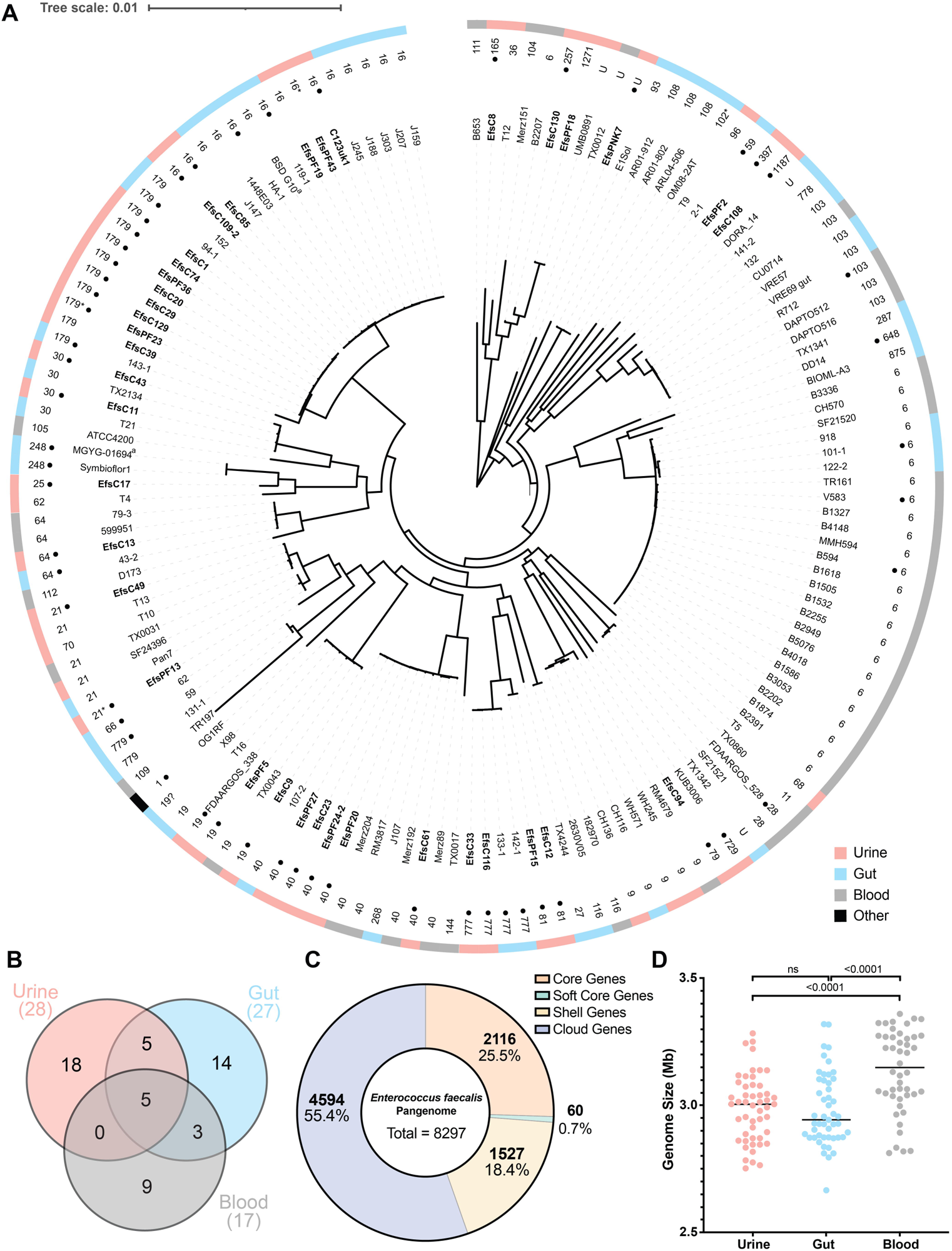
Phylogenetics, MLST, pangenome and genome size distributions of urinary, gut, and blood *E. faecalis* isolates. **(A)** MinVAR-rooted maximum likelihood phylogenetic tree constructed from the core gene alignment of all strains in this study. Isolate names are listed at leaves. Dots represent complete genome assembly. ST is depicted next to isolate names. U: unknown, *: ST has a novel allele ?: ST uncertain. Outermost ring is color-coded by isolation source: urine (pink), gut (blue), blood (grey), reference (black). **(B)** MLST Venn diagram depicting the total number of distinct sequence types in each of the isolation groups. Totals are listed below group names. **(C)** Pangenome analysis summary. Core genes present in >99% of isolates, Soft Core genes present in 95-99% of isolates, Shell genes present in 15-95% of isolates, and Cloud genes present in <15% of isolates. **(D)** Distributions of genome size in megabases of isolates per isolation group. Isolates represented by dots and the line represents median. Statistical significance was determined using ordinary one-way ANOVA with multiple comparisons post-hoc. ^a^Isolate name has been shortened for simplicity. MGYG-01694 - MGYG-HGUT-01694, BSD G10 - BSD2780061688st3_G10.

Pangenome analysis indicated that *E. faecalis* has an open pangenome consisting of a large proportion of accessory genes (soft core, shell, and cloud) and small proportion of core genes (25.5%). In total, 8297 genes make up the pangenome (**Figure 2C**). Isolation source-specific pangenome analysis reveals a similar trend further emphasizing the diversity among urinary isolates (**Figure S1**). Analysis of genome size distribution showed that genomes from blood isolates are on average significantly larger than urine and gut (**Figure 2D**). Conversely, no significant difference in average genome size was detected between urine and gut isolates. These data indicate that urinary isolates are evolutionarily diverse encompassing many STs and a wide range of genome sizes. Furthermore, these findings support the hypothesis that urine isolates likely originate from gut reservoirs (14, 15, 19, 35).

### Plasmid replicon (rep) typing suggests similarities in plasmid carriage between urinary and gut isolates

Plasmids are drivers of evolution and virulence in bacteria (36). We hypothesized that urinary isolates possess characteristic conserved plasmid replicon (rep) types due to within-niche specialization. PlasmidFinder identified 18 unique rep types among all *E. faecalis* strains, 8 of which were present in all groups (repUS43, rep9b, rep9a, rep9c, repUS11, rep2, rep1, rep7a) (**Figure 3A, Table S4**). An extrachromosomal element within EfsPF36 was not typeable by PlasmidFinder or NCBI PGAP. Prophage analysis determined the extrachromosomal element, named here EfsPF36_phage01, is an intact prophage, phiFL2A (**Table S4**).

**Figure 3.**
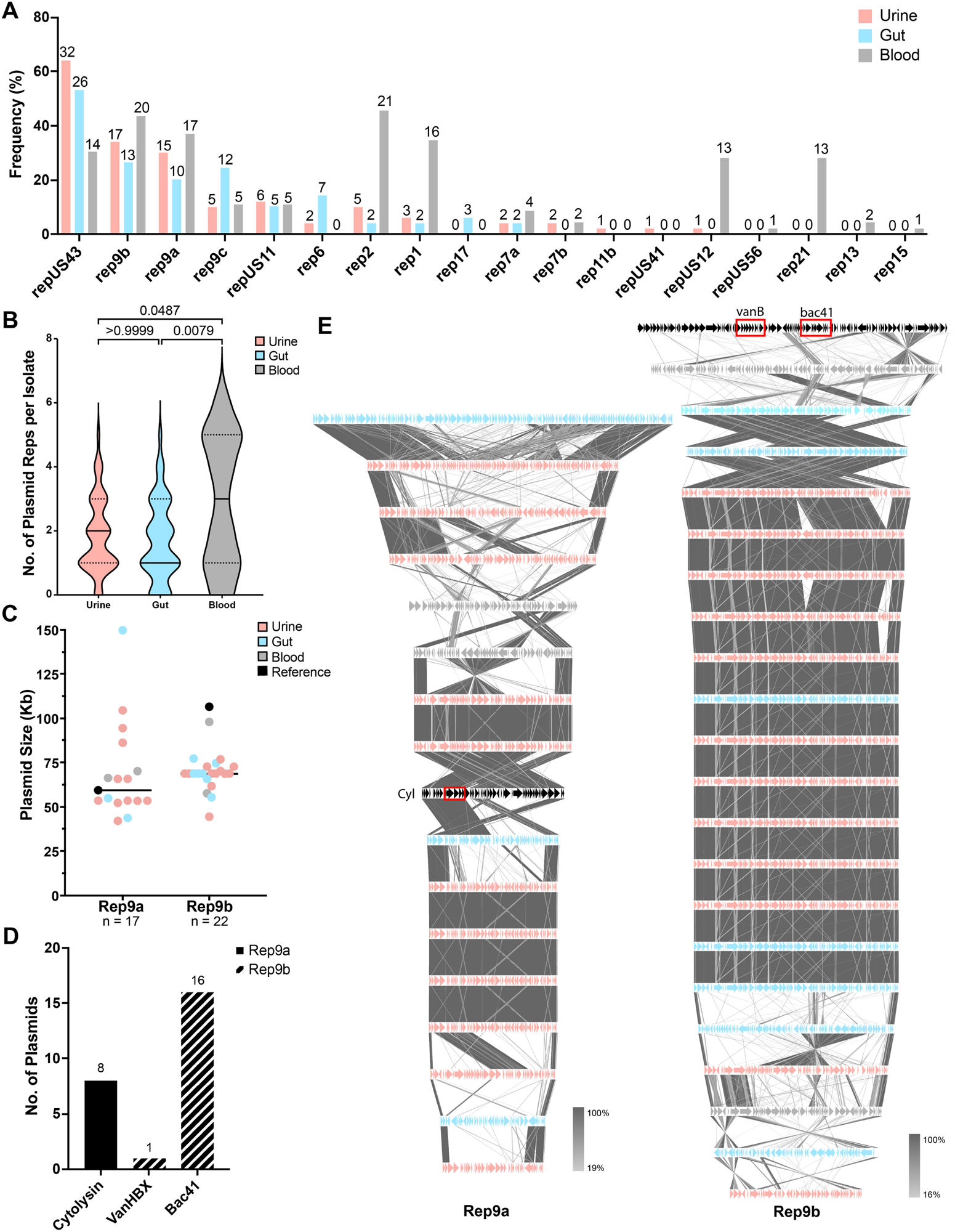
Replicon (rep) type analysis and in-depth comparison of rep9a and rep9b. **(A)** Frequency (raw counts normalized to total group size) of all reps identified. Isolate counts are listed at bar top. **(B)** Distribution of number of plasmid reps per isolate in each isolation group. Statistical significance was determined using Kruskal-Wallis test with multiple comparisons. **(C)** Size in kilobases of each rep9a and rep9b plasmid within complete genome assemblies. Each dot represents a single plasmid and is color-coded by isolation group. Rep9a n=17, Rep9b n=22. **(D)** Number of plasmids possessing cytolysin, vancomycin resistance operon VanHBX, and bacteriocin Bac41 as identified by sequence alignments. Bars are colored by the plasmid rep type associated with the locus. **(E)** tblastx alignments of rep9a (left, n=17) and rep9b (right, n=22) complete plasmids. Arrows denote coding sequences and are color coded by plasmid isolation source: urine (pink), gut (blue), blood (grey), reference (black). Reference rep9a plasmid is DS16 pAD1. Reference rep9b plasmid is pMG2200. Shaded lines between plasmid sequences are colored based on sequence identity (%). Locus of cytolysin operon, VanHBX operon, and bac41 are highlighted in the reference plasmid by red blocks.

Rep6 was the only rep present in urine and gut isolates, but not in blood. Rep7b and repUS12 were only present in urine and blood isolates but not in gut. Of the rep types that were unique to a single isolation group, rep17 was only identified among gut isolates, rep11b and repUS42 were identified only in urine isolates, and repUS56, rep21, rep13, and rep15 were only identified in blood isolates (**Figure 3A**). We found that the blood group possessed the total highest number of replicons (n=134) followed by urine (n=92) and gut (n=82) (**Figure 3A**). Additionally, the blood group had the highest median number of replicons per isolate compared to urine and gut (**Figure 3B**). Urinary and gut isolates appeared closely related in terms of plasmid types present in these strains while blood isolates possessed more unique rep types.

The most common rep type was repUS43 found in 64% of urine, 53% of gut, and 30% of blood isolates (**Figure 3A**). Analysis of the complete, closed genomes generated in this study revealed that repUS43 is chromosomally integrated and is proximal to a tetracycline resistance gene (*tetM)* (**Table S4**). The following two most prevalent replicons were rep9a and rep9b which are associated with pheromone responsive plasmids. rep9a is associated with pAD1 lineage plasmids known to encode the virulence factor cytolysin (37–39). Additionally, rep9b is associated with well-characterized pTEF2 of model strain V583, reported to encode vancomycin resistance (*vanHBX*) and a Bac41 type bacteriocin (40). We found that identified rep9a plasmids were diverse in size, ranging from 149 kb to 42 kb and only 47% encoded the cytolysin operon (**Figure 3C-D**). Alignments of complete rep9a plasmids and the DS16 pAD1 reference further suggests their genomic content varies widely (**Figure 3E**). Rep9b plasmids also have a broad size range from 106kb to 44kb, however, 40.9% of complete rep9b plasmids are approximately 68kb (**Figure 3C**). Alignments of all complete rep9b plasmids and the pMG2200 reference indicates that *vanHBX* is only encoded in pMG2200 while 72.7% of rep9b plasmids encoded the *bac41* operon (**Figure 3D-E**).

### Pseudolysogenic extrachromosomal phage is enriched among geographically proximal isolates

Complete genome assemblies of the urinary collection revealed the presence of a prevalent pseudolysogenic extrachromosomal phage, EF62phi (41). The phage, EF62phi, originally identified in *E. faecalis* 62 (SAMN02603509) isolated in Norway, encodes common structural and phage components as well as a toxin-antitoxin system (**Figure 4A**). EF62phi was identified in 13 (26%) of the urinary isolates sequenced as part of this study. The phage was also identified in 18.4% of gut and 8.7% blood isolates (**Figure 4B, Table S4**). Notably, EF62phi was commonly found in gut strains isolated in the same geographical region as the urinary collection. Thus, we hypothesized that EF62phi was associated with isolate geographical origin. Analysis utilizing available geographical metadata revealed that a greater percentage (36.5%) of *E. faecalis* isolated from the Dallas Metroplex in Texas, USA harbored EF62phi than Non-Dallas isolates (8.97%) (**Figure 4C**).

**Figure 4.**
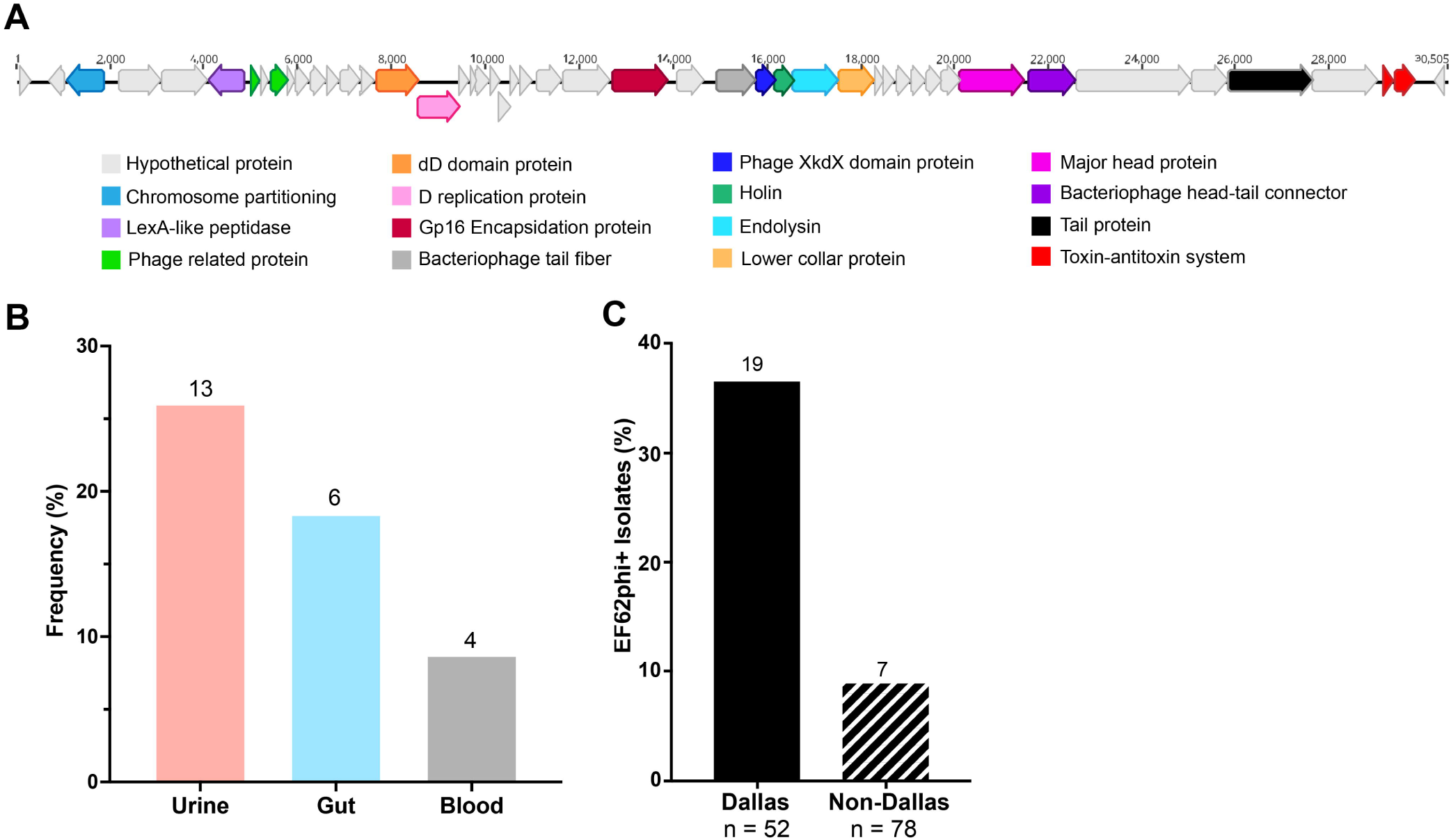
Extrachromosomal pseudolysogenic bacteriophage EF62phi prevalence in *E. faecalis*. **(A)** Complete genome of EF62phi with color coded annotation of coding sequences. **(B)** Frequency of presence of EF62phi within genomes of each isolation group. Raw isolate counts are listed at bar tops. **(C)** Number of strains possessing EF62phi that were isolated from the Dallas (n=52) versus non-Dallas (n=78) areas. Raw isolate counts are listed at bar tops. Geographies were determined for a subset of genomes for which geographical metadata was available.

### Antimicrobial resistance is not widespread among urinary E. faecalis isolates

Antimicrobial resistance genes (ARG) analysis was used to predict *E. faecalis* resistance and assess differences between the isolation groups. We hypothesized that urinary isolates would be commonly resistant to UTI front-line therapies such as nitrofurantoin and fluoroquinolones. Strikingly, the median number of ARGs, including known point mutations, per isolate was lowest in the urine group (**Figure 5A**). Resistance was predicted to 11 drug classes (tetracycline, macrolide, aminoglycoside, trimethoprim, vancomycin, phenicol, beta-lactam, lincosamide, Fosfomycin, nitrofurantoin, and fluoroquinolone) attributed to 31 unique ARGs or mutations in *E. faecalis* from all three isolation groups (**Figure 5B, Table S5**). Several ubiquitous genes confer intrinsic resistance to diverse antimicrobials in the species. These genes include *lsaA*, *lsaB*, *lsaE* encoding multidrug resistance ABC-F subfamily efflux pumps, *dfrE* encoding a dihydrofolate reductase responsible for trimethoprim resistance, *efrA* and *efrB* encoding components of the EfrAB efflux pump conferring multidrug resistance, and *emeA,* encoding another efflux pump (**Table S5**) (42–46). The most prevalent tetracycline resistance gene was *tetM,* with gut isolates most frequently and blood isolates least frequently encoding tetracycline resistance (67% gut, 56% urine, 46% blood). The *tetM* gene was commonly associated with the chromosomally integrated repUS43 locus and the prevalence of repUS43 correlated with the presence of *tetM*, (Spearman r=0.7271, *p*<0.0001) (**Figure 5B**). Intriguingly, tetracycline was the only drug class for which urinary isolates were more frequently predicted to possess ARGs than blood isolates.

**Figure 5.**
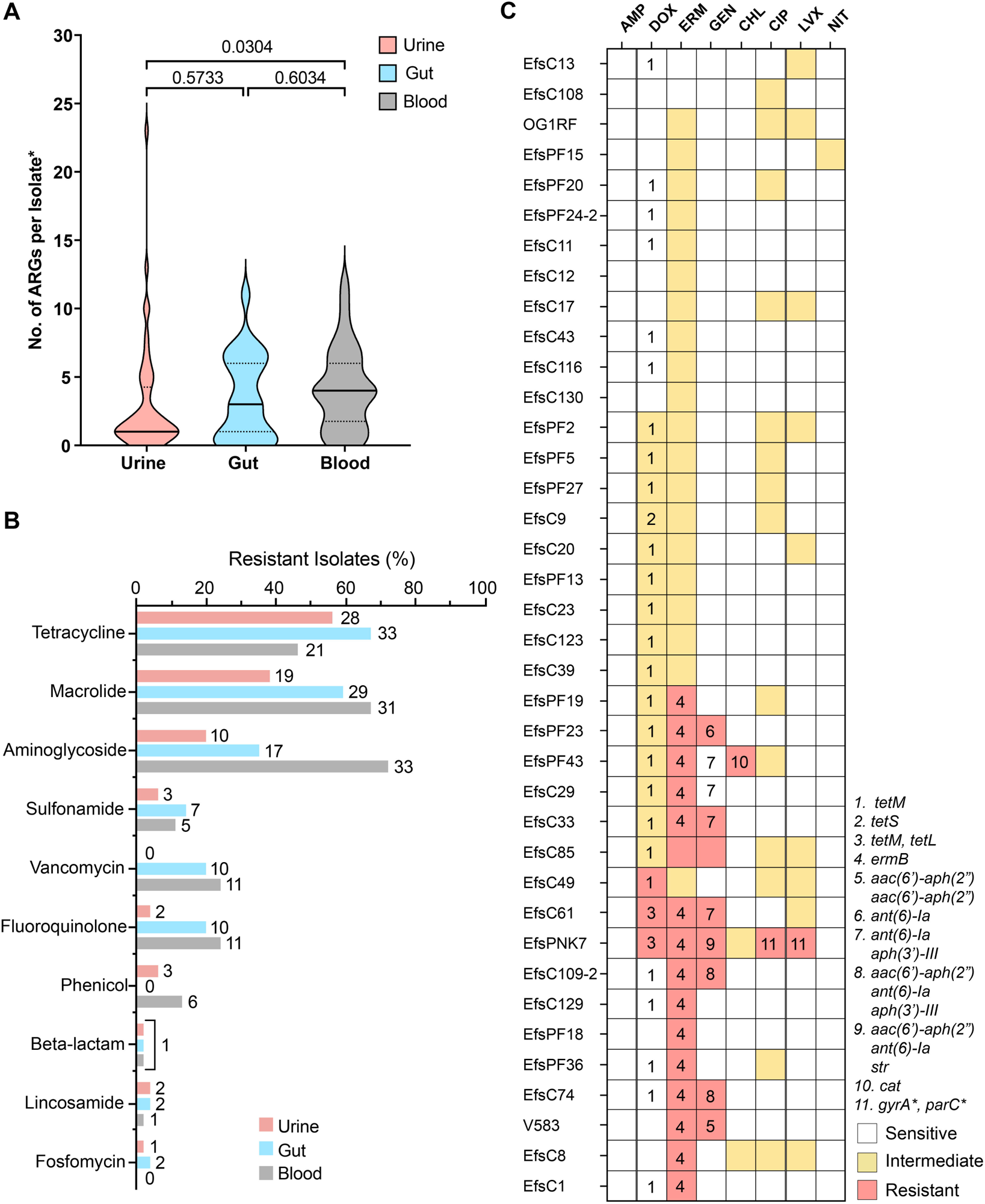
Antimicrobial resistance genes and chromosomal mutation predictions and resistance phenotypes of urinary isolates. **(A)** Distribution of total number of ARGs per strain in each isolation group. Each copy of a multi-copy ARG was counted. *Chromosomal mutations were counted as ARGs. Statistical significance was determined using Kruskal-Wallis test with multiple comparisons. **(B)** Frequency of resistant isolates in each isolation source as predicted by ARG in silico analysis. Isolate counts are listed as bar ends. **(C)** Heatmap of resistance phenotypes assessed by disk diffusion and MIC assays. Numbers correspond to presence of an ARG or mutation. Ubiquitous efflux pumps and intrinsic resistance genes are not depicted.

Aminoglycosides, fluoroquinolones, and vancomycin demonstrated substantial differences in prevalence between the three isolation groups (**Figure 5B**). In all three cases, urine isolates were predicted least frequently to be resistant whereas blood isolates were predicted to be most frequently resistant. This in part may be due to bias in the selection criteria of sequenced *E. faecalis* blood strains – most from hospital outbreaks, surveillance, et cetera (24, 33) – that target Vancomycin-resistant and MDR strains. Aminoglycoside resistance was substantially more common in blood isolates (72%) compared to gut (35%) and urine (20%) strains (**Figure 5B**). Fluoroquinolone resistance, as predicted by the presence of either an ARG or known *gyrA* and/or *parC* mutations, was similarly prevalent in gut (20%) and blood (24%) isolates, but less frequent in urine isolates (4%) (**Figure 5B**). Because fluoroquinolone resistance is used less frequently as selection criteria for *E. faecalis* strain isolation, this finding may be more representative of mutation frequencies of *E. faecalis* in gut, blood, and the urinary tract. Finally, vancomycin resistance was entirely absent in the urine group but found in similar frequency in gut (20%) and blood (24%) isolates (**Figure 5B**).

Although gene predictions offer insight into possible resistance phenotypes, phenotypic assessments are imperative to confirm resistance. Phenotypic antibiotic resistance analysis of a large collection of urinary isolates has not been previously conducted, therefore we analyzed resistance phenotypes of 38 strains to 8 clinically relevant antibiotics from 7 distinct drug classes (**Figure 5C**). OG1RF and V583 were included as controls and representatives of well-studied *E. faecalis* model strains. Resistant phenotypes among urinary isolates largely converged with gene prediction data where specific resistance genes were present, whereas ubiquitous efflux pumps were not associated with resistance in all strains (**Figure 5C, Table S5**).

Thirty resistant phenotypes were observed, of which 29 (96.6%) were attributed to the presence of a predicted ARG or known chromosomal mutations (**Figure 5C**). Resistant phenotypes to erythromycin were most widespread as expected from gene prediction with 42% of isolates demonstrating resistance (**Figure 5C**). Erythromycin resistance was well-predicted by the *ermB* gene – of 16 erythromycin resistant isolates, 15 possessed *ermB*. Gentamicin resistance was the second most prevalent with 8 resistant isolates. Gene prediction analysis in correlation with the resistance phenotypes suggests that *aac(6’)-aph(2”)* may primarily be responsible for resistance, with 5 of 8 resistant isolates encoding this gene (**Figure 5C**). Gentamicin resistance in EfsC85 was not explained by gene prediction analysis and the mechanism for resistance in this isolate remains unclear (**Figure 5C**).

The third most prevalent resistance was to doxycycline of the tetracycline class. The presence of *tetM* gene, the most common of tetracycline ARGs, was not uniquely associated with resistance. In isolates encoding only *tetM*, one isolate was resistant, 14 were intermediate, and 11 were susceptible. Co-occurrence of *tetM* and *tetL* was a more accurate predictor of doxycycline resistance (**Figure 5C**). Chloramphenicol resistance was only predicted in one isolate and was associated with the presence of the *cat* gene. Finally, only one isolate was resistant to ciprofloxacin and levofloxacin resistance due to presence of known mutations in the QRDR regions of the *gyrA* and *parC* genes (47) (**Figure 5C**).

Beyond resistant phenotypes, a total 61 intermediate phenotypes were observed, of which 15 (39.4%) were attributed to the presence of a predicted ARG (**Figure 5C**). The remaining 46 intermediate phenotypes were largely detected for erythromycin (20), fluoroquinolones (ciprofloxacin, 14; levofloxacin, 9) and less commonly for chloramphenicol (2) and nitrofurantoin (1). Because of their clinical relevance, we further analyzed intermediate resistance to fluoroquinolones. We first validated ciprofloxacin intermediate resistance phenotypes by MIC assay (**Table S6**) and confirmed that reported intermediate strains had ciprofloxacin MICs ≥2 and <4μg/mL. Interestingly, no known point mutations or fluoroquinolone resistance ARGs were associated with the validated intermediate phenotypes (**Figure 5C**).

### Genes involved in sugar transport and metabolism, vitamin B12 import, and post-transcriptional stress responses are enriched in urinary E. faecalis

We next sought to identify potential mechanisms of genetic adaption to the urinary niche by evaluating gene enrichments associated with urinary versus gut strains of *E. faecalis.* Following inclusion cutoffs of >70% frequency in urine isolates and ≥20% higher prevalence in urine than in the comparator group, gene enrichment analysis identified 19 candidate genes as enriched in urinary isolates when compared to gut (**Figure 6A, Table S7**). Among the 19 genes, 6 encode a PTS system operon predicted to be responsible for the transport of mannose and fructose (*manR*, *fryA*, *manP_1*, *manP_2*, *fruA*, *alsE*). Eight genes are located within a syntenic region of prophage phage 04 of V583 (locus EF1988 – EF2043) (48, 49). These 8 candidates include 4 hypothetical proteins (group_887, group_1388, group_2882, group_3220); 2 phage components, ArpU family transcriptional regulator and a recombinase (*xerC*); and 2 intriguing, annotated candidates: *csp* encoding a cold-shock protein and *hemH* encoding a ferrochelatase (**Figure 6A, 6B**). *Csp* is a unique member of the cold shock protein (csp) family. There are 7 *csp* orthologs in the *E. faecalis* pangenome, 4 of which are present in all *E. faecalis* strains. Among the *csp* genes that are not ubiquitous, the enriched *csp* candidate (EF1991) was found to be a unique cold shock family member present at higher frequency (86%) within urinary isolates (**Figure S3**). The location of these genes within a prophage region suggests they may have been acquired by horizontal gene transfer through a phage integration event. To determine if the enrichment of these 8 prophage-associated candidate genes was specific and not just a result of enrichment of the entire prophage region, we aligned the phage region sequenced and performed blastn queries of candidate genes (**Figure 6C**). These data along with the presence/absence data suggest that the candidate genes are enriched independently of the intact phage because the phage structural genes, for example, are found at much lower frequencies among urine isolates than the 8 enriched genes (**Figure 6C, Table S7**).

**Figure 6.**
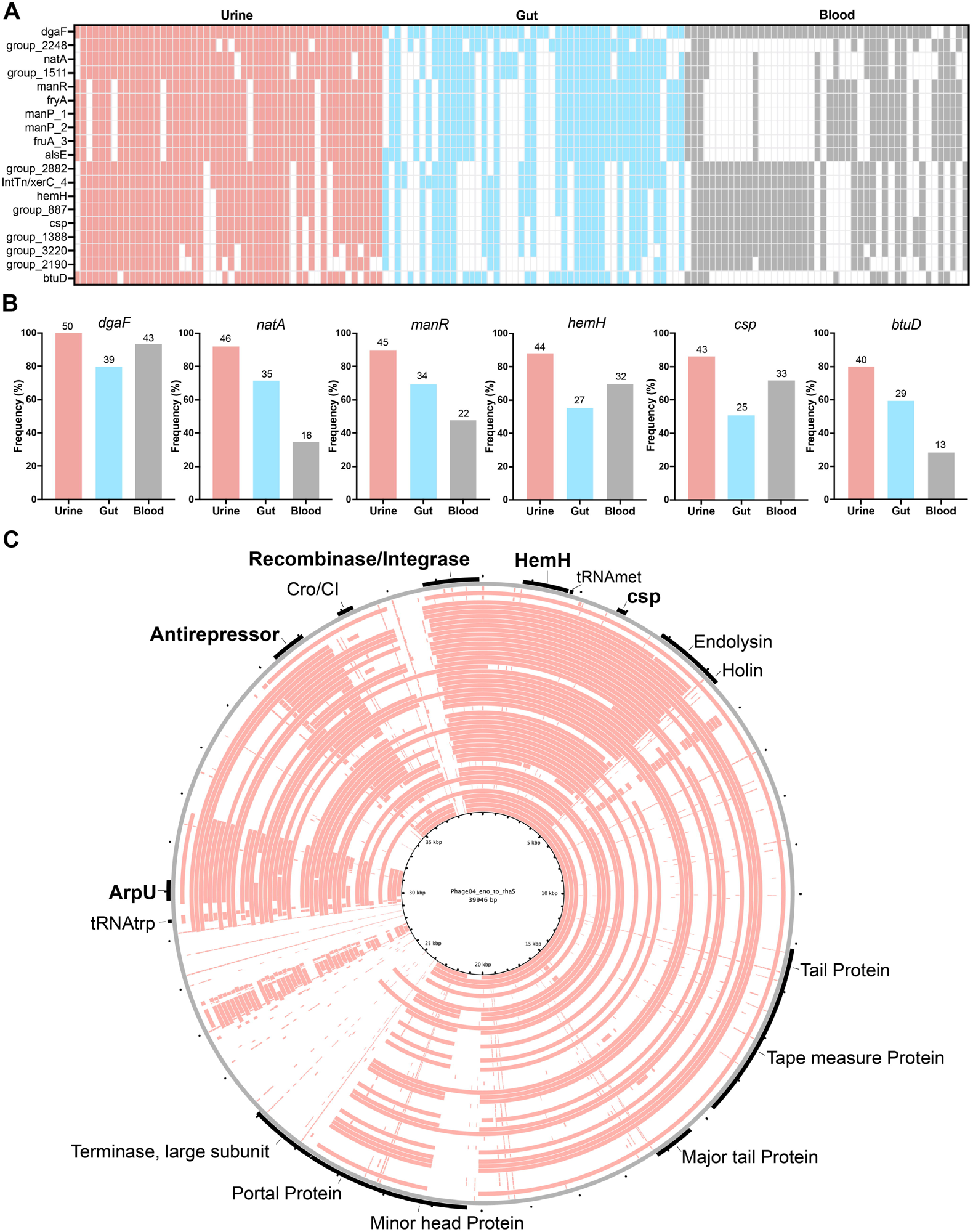
Candidate genes enriched among urinary *E. faecalis* isolates. **(A)** Presence/absence map of 19 candidate genes. Columns represent a single isolate, colored blocks correspond to isolation source: urine (pink), gut (blue), blood (grey). Genes were identified comparing urine and gut isolates, presence in blood isolates is provided for reference. **(B)** Frequency of candidate genes within each isolation source. Isolate counts are listed at bar tops. **(C)** BLAST Ring Image Generator (BRIG) blastn alignment of urinary isolates (pink rings) phage04 integration region (*eno* to *rhaS*) to V583 phage04 reference (grey ring). Prophage annotations are listed and enriched candidates are in bold.

The remaining 5 enrichment candidates include *dgaF*, *natA*, *btuD*, and two hypothetical proteins (group_1511, group_2248) one of which is directly downstream of *natA* (**Figure 6A, 6B**). DgaF is an aldehyde lyase used in the Entner-Doudoroff pathway that *E. faecalis* is known to employ in its carbohydrate metabolism (50, 51). NatA is a sodium ABC transporter, ATP binding protein while BtuD is a vitamin B12 import ATP binding protein. In the *E. faecalis* pangenome there are up to 19 distinct *btuD* orthologs, some present in all isolates. A single *E. faecalis* may possess between 6 to 10 of the *btuD* orthologs ranging in length from 768bp to 2340bp. The enriched *btuD* candidate encodes a 297aa protein. Its length as well as sequence composition suggests it is a unique family member of the vitamin B12 importer family (**Figure S4**).

All hypothetical candidates were analyzed using NCBI conserved domains database, and none were found to have conserved domains. However, blastp analysis found that candidate group_2248 was 100% identical to a T7SS effector LXG polymorphic toxin (accessions CP091889, WP_238463980.1) and group_1511 downstream of *natA* was 99.1% identical to an annotated ABC transporter, permease protein (accession HF558530). This coupled with the respective lengths of *natA* and group_1511 suggest that the latter is *natB* (52). Candidate group_2882 of the phage04 region was 100% identical to an anti-repressor (accession CP014949) (**Table S7**).

## METHODS

### Urinary E. faecalis Strain Isolation

Clean catch midstream urine specimens were collected from female patients following informed patient consent and institutional review board (IRB) approval (STU 082010-016, STU 032016-006, MR 17-120). Patients were stratified into four groups based on their UTI history, symptoms, and urinalysis results at the time of collection: Never, Sporadic, rUTI Remission, and rUTI Relapse. Here, rUTI is defined as ≥2 symptomatic UTIs within six months or ≥3 UTIs within 12 months (17, 53, 54). Patient age was >35 with majority being postmenopausal (>55 years old). Additional clinical metadata including Body Mass Index (BMI), smoking history, diabetes status, electrofulguration of trigonitis history, and prescription of estrogen hormone therapy (EHT), non-steroidal anti-inflammatory drugs (NSAIDs), and antibiotics were recorded for analysis (**Table S2**). Urinary *E. faecalis* isolates were cultivated from urine specimens and preliminarily identified by plating 100 or 10 µL of urine on CHROMagar Orientation and incubating at 37°C overnight. Species identity was confirmed for isolates exhibiting *E. faecalis* colony morphology by PCR of the 16S rRNA gene and Sanger sequencing, followed by megablast query at default thresholds as described previously (55–58).

### Urinary, gut and blood E. faecalis strain genome collection

A-priori power analyses using one-way ANOVA and Fisher’s Exact Test (G*Power) were conducted to determine sample size for the comparative analysis of two or three groups (59). This analysis determined that to detect an effect size of 0.4 with a power of 0.8, a minimum of 41 genomes are needed in each isolation group. A total of 146 *E. faecalis* strains were included in this study: 50 urine isolates, 49 gut isolates, 46 blood isolates, and OG1RF, a commonly studied oral commensal *E. faecalis* (**Table S3**). Among the 49 gut isolates, 15 genomes of *E. faecalis* gut isolates obtained from patients in Dallas, Texas, USA area hospitals were included as they represent geographically relevant comparator strains from the human GIT and are high quality closed genomes (BioProject PRJNA800580). *E. faecalis* genome assemblies available as of July 28, 2020 were downloaded from the NCBI assembly database following stringent curation. Inclusion criteria were as follows: human host, isolated from the gastrointestinal tract (gut, fecal), blood, or urine; isolated from distinct individuals (non-longitudinal); meet assembly quality parameters of average coverage >30x, contig N50 >80,000 and fewer than 100 contigs. Finally, isolates from published research were prioritized.

### Genomic DNA Isolation and Sequencing

Genomic DNA (gDNA) was extracted using Qiagen Blood & Tissue kit and assessed for quality using 260/280nm absorbance ratio, fluorometry, and agarose gel electrophoresis. High quality gDNA libraries were prepared for short-read Illumina and long-read MinION sequencing as described previously (56, 60). See detailed methods in **Supplemental Materials**.

### Genome Annotation & Sequence Type Predictions

Genome annotation was performed using Prokka v1.14.6 and NCBI Prokaryotic Genome Annotation Pipeline v6.4 with default parameters (61, 62). Multi-locus sequence typing was performed using MLST v2.0 on the Center for Genomic Epidemiology (CGE) server with the *E. faecalis* configuration and Assembled or Draft Genome/Contigs as input (63, 64).

### Plasmid Replicon Analysis

Plasmid replicon types were predicted using PlasmidFinder v2.1 at default parameters (64–66). Extrachromosomal elements not typeable by PlasmidFinder were assessed using NCBI Conserved Domain Search at default parameters, PHASTER, and blastn queries of the nr/nt database (67–69). PLSDB v2021_06_23_v2 was used with both Mash screen and Mash dist search strategies to identify similar plasmids (70, 71). Alignment of complete plasmid sequences was conducted using Easyfig v2.2.2 with tblastx at default parameters (72). Reference plasmids representing rep9a (pAD1 from strain DS16, SAMN00809239) and rep9b (pMG2200, AB374546.1) were included in the analysis for comparison. Further alignment and visualization were conducted using BLAST Ring Image Generator (BRIG) v0.95 with blastn at default thresholds.

### Pangenome & Phylogenetic Analysis

Pangenome analysis was conducted using Panaroo v1.2.10 with merge-paralog option selected and core gene alignment conducted using mafft (73, 74). Core gene alignment output was then used to construct a phylogenetic tree with IQ-TREE using the GTR+F+I+I+R6 model with ultrarapid bootstrapping (1000 inferences) (75–77). Model of nucleotide substitution was selected with IQ-TREE model selection (77). The phylogenetic tree was rooted using min-VAR rooting with FastRoot v1.5 and visualized with iTOL (78, 79).

### Gene Enrichment Analysis and Functional Annotation

Gene enrichment analysis was conducted using Scoary v1.6.16 (80). Genes with p-value <0.05 were retained for further analysis (**Table S7**). Candidates were considered enriched if they were present in >70% of urine isolates and were at least 20% more prevalent in urine isolates than in the comparator group. Since the urine group is primarily comprised of isolates sequenced in this study (74%), it was imperative to focus on candidates present in more than 70% of isolates to ensure biological significance. Candidates were validated using blastn to confirm paralogs were not erroneously split and ensure highly conserved homologs are not unaccounted for. Hypothetical candidate genes were further assessed using NCBI Conserved Domain Search, PSORTb v3.0.3, blastn and blastp queries to elucidate potential function (68, 69, 81). Allele sequence alignments were conducted using nucleotide MUSCLE v3.8.425 alignment at default parameters in Geneious prime v2022.1.1 (82, 83).

### Antimicrobial Resistance Gene Prediction

Antimicrobial resistance genes and point mutations were predicted using ResFinder v4.0 on CGE with the *E. faecalis* database at default parameters (84, 85). ABRicate v1.0.1 was used to query the CARD database at 70% coverage and 70% identity thresholds to validate ResFinder findings and identify additional resistance determinants (86, 87).

### Antimicrobial Resistance Phenotype Assessment

Resistance phenotypes were assessed by Kirby-Bauer disk diffusion, as described previously, on Brain-Heart Infusion (BHI) agar and antimicrobial susceptibility was evaluated by measurement of the zone of inhibition per the established zone diameter breakpoints of Clinical and Laboratory Standards Institute (CLSI) (88–90). All Ciprofloxacin phenotypes and intermediate or resistant Gentamicin and Chloramphenicol phenotypes were further validated using Minimum Inhibitory Concentration (MIC) microdilution assay in BHI broth per CLSI breakpoints. MIC was measured using the HT-MIC workflow as previously described (91). EfsC94 of the urinary collection was excluded due to insufficient growth. Further details can be found in **Supplemental Materials**.

## DISCUSSION

In this study we analyze the genomes of urinary, gut, and blood *E. faecalis* isolates with the purpose of elucidating genetic adaptations of *E. faecalis* to the urinary tract. A collection of 37 clinical urinary isolates was obtained from predominately postmenopausal women, sequenced, and hybrid-assembled to generate closed or highly contiguous genome assemblies. This collection, first and foremost, provides a tool for further research in the fields of *E. faecalis* genomics and *E. faecalis* UTI. Coupled with additional curated publicly available *E. faecalis* genomes, this collection also offers a comprehensive insight into population genetics, mobile genetic elements, antimicrobial resistance, and enriched genes that may help distinguish urinary *E. faecalis* from strains of other human anatomical niches.

Here, we sought to test the hypothesis that urinary *E. faecalis* possess genetic adaptations that enable colonization of the urinary environment. We postulated that urinary isolates should be genetically conserved, hold a characteristic plasmid content, be resistant to similar antimicrobials, and encode genes that may aid in survival within the urinary tract. Additionally, as the gut-bladder axis becomes further defined in UTI research, we estimated that urinary isolates will resemble gut isolates more than blood isolates (15, 19, 35).

Our findings reveal that urinary *E. faecalis* is diverse and no lineage emerges as strongly associated with the urinary niche. The most frequent ST in the urinary group, ST179, is a single locus variant of the most common ST in the gut group, ST16. Strains belonging to these two STs form a large phylogenetic cluster and may support the gastrointestinal origin of urinary isolates. Blood isolates demonstrate an expected lineage bias for ST6 (24). As genome availability is often a reflection of study selection criteria, ST6 blood isolates have been primarily associated with hospital outbreaks and represent a large collection of strains available publicly for analysis (24). We recognize this introduces inevitable bias to the analysis and cautiously draw conclusions about urinary versus blood comparisons.

Plasmid replicon analysis did not identify strong associations between rep type and isolation source. With exception of two rep types (rep11b, repUS41) which are both present in a single isolate, EfsC94, no rep types were found to be strictly associated with urinary isolates. Most rep types were represented in all three isolation groups with pheromone responsive rep9a, rep9b, and rep9c being among the most prevalent. RepUS43 was the most common rep type and was found to be chromosomally integrated within all complete genome assemblies assessed in this work. Furthermore, repUS43 was positively correlated with the presence of tetracycline resistance gene *tetM* suggesting this resistance gene may be widespread due to a recombination event with a repUS43 plasmid in a common ancestor. Rep typing offered an insight into the plasmid content of urinary *E. faecalis* but not without limitations. As the analysis focuses on a single locus, the remaining genomic content of the associated plasmid remains unclear. Alignments of complete rep9a and rep9b plasmids revealed that in both cases, plasmids of these rep types can vary widely in terms of size as well as possession of significant traits such as cytolysin, vancomycin resistance, and bacteriocin. Rep typing thus may be more useful for specific replicons but should be utilized with caution when considering plasmid function.

Antimicrobial resistance *in silico* predictions indicated that urinary *E. faecalis* encode less ARGs than gut or blood counterparts. In all drug classes for which ARGs were detected in the genomes of isolates, with two exceptions, urinary isolates possessed ARGs less frequently than both gut and blood isolates. The exceptions were the tetracycline class, in which tetracycline ARGs were more prevalent in urine than in blood isolates and the beta-lactam class for which only a single isolate from each niche was found to be resistant. Acquired ARGs or mutations predicted to confer fluoroquinolone or nitrofurantoin resistance were not prevalent in urine isolates. Phenotypic *in vitro* assessments of antibiotic susceptibility of urinary isolates demonstrated a similar trend by which resistance was not widespread and intermediate phenotypes were more prevalent. Intermediate fluoroquinolone resistance was relatively common, but the mechanism remains unclear. Uncharacterized *gyrA* and *parC* mutations were present in some strains, albeit not within the QRDR regions, but may partially explain the observed intermediate phenotypes. A limitation of this study is that the strains used in this analysis were isolated from the same clinic and are likely not fully representative of diverse isolation sites, clinical practices, and demographic groups.

To address the hypothesis that urinary *E. faecalis* possesses genetic and phenotypic adaptations that enable its survival in the urinary tract, we conducted gene enrichment analysis comparing urinary and gut isolates. We focused on these two isolation groups to identify genes that may play a role in the adaptation of *E. faecalis* from the gut to the urinary tract. Applying stringent inclusion cutoffs, we identified 19 candidate genes enriched in the urinary group. Of note, we found an enrichment of genes encoding a mannose/fructose PTS system. This may be relevant to the urinary niche because the glucose concentration in urine is often low (0.2-0.6mM), and therefore this operon may help *E. faecalis* utilize alternative carbon sources that are more abundant in urine like mannose (28). Intriguingly, D-mannose supplements are routinely advised for UTI prophylaxis as studies suggest it may block colonization by uropathogenic *E. coli* which adhere to the host urothelium via mannosylated cell surface proteins (92, 93).

Additional candidates like *dgaF* which functions as a 2-keto-3-deoxygluconate 6-phosphate (KDGP) aldolase to form glyceraldehyde-3-phosphate and pyruvate similar to the Eda enzyme in the Entner-Doudoroff (ED) pathway and is found in all urinary isolates may be important in *E. faecalis* growth under the environmental pressures of the urinary niche (94). The ED pathway produces less ATP than glycolysis but offsets this metabolic cost by requiring less enzymes. While the ED pathway is often used by aerobic Gram-negative bacteria, *E. faecalis* is one of the few facultative anaerobic Gram-positive microbes that possess it (50, 95). We postulate that *E. faecalis* may utilize the ED pathway during urinary colonization to conserve protein and therefore energy costs (96).

Another gene observed to be enriched among urinary strains was *btuD*. This gene encodes an ABC ATP binding protein predicted to be involved in cobalamin (vitamin B12) transport (97). Cobalamin is an important cofactor in the ability of bacteria to utilize ethanolamine as a carbon source (98). In fact, ethanolamine, which is relatively abundant in urine, has been proposed to provide a growth advantage for UPEC by offering a carbon and nitrogen source for rapid colonization of the urinary tract (99). Murine models also suggest ethanolamine metabolism is essential for bladder colonization of UPEC (100). It is conceivable that cobalamin transport mediates ethanolamine utilization in *E. faecalis*.

Lastly, two enrichment candidates of a close genomic localization with a prophage region were *hemH* and *csp*. HemH is a ferrochelatase known to insert ferrous iron to form coproheme in Gram-positive bacteria (101). However, *E. faecalis* lacks the machinery to synthesize heme and thus the role of this ferrochelatase remains unclear (102). Of particular interest is *csp* encoding a cold shock protein (CSP). CSPs are commonly encoded by bacteria as an adaptive mechanism to stress and changes in environmental conditions. The first CSP, CspA, was identified in *E. coli* and later found to have nine homologs (CspA – CspI). These proteins, commonly expressed in response to temperature decreases, function as chaperones that prevent the formation of secondary structures in RNA transcripts thereby allowing proper translation (103, 104). However, some CSPs are known to be non-cold inducible. In *E. coli* four of the nine CspA family members are not induced by cold shock. Such CSPs have been reported to be involved in osmotic, oxidative, starvation, pH and ethanol stress tolerance as well as host cell invasion(103, 104). An example of such adaptive CSP has been described by Michaux *et al*. (104) which suggested CspR of *E. faecalis* plays a special role in virulence and persistence (104). The enriched *csp* gene highlighted by our analysis was shown in previously to be upregulated in V583 grown in blood (27). However, the function of the CSP allele enriched in urinary isolates and its role in urinary fitness has yet to be characterized in *E. faecalis*.

## ACKNOWLEDGEMENTS

We thank The University of Texas at Dallas Genome Center for their services and expertise. This work was funded by the Welch Foundation, award number AT-2030-20200401 to N.J.D., by the National Institutes of Health, award number R01AI116610 and the Cecil H. and Ida Green Chair in Systems Biology Science to K.P., and by the Felecia and John Cain Distinguished Chair in Women’s Health, held by P.E.Z.

## AUTHOR CONTRIBUTIONS

Conceptualization: N.J.D., K.L.P., B.M.S., Data Curation: B.M.S., A.P.A, A.N., S.T., S.S.R.B., N.V.H., D.P.A. Formal analysis: B.M.S., A.P.A, A.N., M.L.N. Funding acquisition: N.J.D., K.L.P., P.E.Z. Investigation: B.M.S., A.P.A, A.N., S.T., S.S.R.B., N.V.H., D.P.A., Methodology: B.M.S., A.P.A, S.T., Project administration: N.J.D., K.L.P., B.M.S., Resources: N.J.D, K.L.P., P.E.Z., N.A.D., Software: B.M.S., Supervision: N.J.D., K.L.P., N.A.D. Validation: N.J.D., K.L.P., B.M.S., Visualization: B.M.S., N.J.D. Writing – original draft: N.J.D., K.L.P., B.M.S., Writing – review & editing: N.J.D., K.L.P., B.M.S., P.E.Z.

## Notes

https://www.ncbi.nlm.nih.gov/bioproject/?term=PRJNA800580

### Competing Interest Statement

The authors have declared no competing interest.

