## Supplementary Material for "Functional and genetic adaptations contributing to *Enterococcus faecalis* persistence in the female urinary tract"

**LIST OF CONTENTS**

Supplemental Methods

Figure S1 - Sequence alignments of complete DS16 pAD1 plasmid with previously published pAD1 sequence fragments

Figure S2 - Pangenomes of urinary, gut, and blood isolation groups

Figure S3 - Nucleotide alignment of all *csp* alleles identified in *E. faecalis*

Figure S4 - Nucleotide identity (%) matrix of all *btuD* alleles identified in *E. faecalis*

**SUPPLEMENTAL METHODS**

***Genomic DNA Isolation and Sequencing***

High quality gDNA libraries were prepared for short-read sequencing using the Nextera DNA Flex library prep kit and sequenced using 2x150bp paired-end sequencing on Illumina NextSeq 500. Long-read sequencing libraries were prepared using Oxford Nanopore Technologies (ONT) ligation sequencing kit (SQK-LSK109) and barcode expansion kits EXP-NBD104 and EXP-NBD114 and sequenced using R9 FLO-MIN106 flow cells on ONT MinION as described previously (32, 36). Sequencing reads were subject to quality filtering and trimming using CLC Genomics Workbench v12.0.3 and NanoFilt v2.6.0 to discard short reads with Phred score <20 and length <15bp, and long reads with Phred score <7 and length <200bp, respectively (37). Genomes were assembled using Unicycler v0.4.8 at default parameters (38-41). Incomplete assemblies were later reassembled using Unicycler v0.5.0 at default parameters or bold mode to obtain the most complete assembly (fewest contigs). Genome completeness was assessed using CheckM v1.0.18 lineage workflow with the Lactobacillales order marker gene set on KBase v1.4.0 (42, 43). Draft genomes contig N50 and quality was assessed using QUAST v5.0.2 (44). All genomes have been deposited to NCBI via BioProject PRJNA944190. Accession numbers are provided in **Supplemental Data S2**.

***Plasmid replicon analysis***

A complete pAD1 plasmid from DS16 was assembled using raw sequence read data of DS16 (SRX158205) using PlasmidSPADES v3.15.2 (49). The complete pAD1 plasmid assembly was validated using global nucleotide alignment with free end-gaps in Geneious Prime v2022.1.1 using publicly available and previously published pAD1 fragments (L01794.1, X62658.1, X62657.1, L19532.1, AB007844.1, X96977.1, X17214.1, AF343839.1, U00681.1) (50-57).

***Antimicrobial Resistance Phenotype Assessment***

Resistance phenotypes were assessed by Kirby-Bauer disk diffusion on Brain-Heart Infusion (BHI) agar to 8 representative antibiotics including: Ampicillin (AMP), Doxycycline (DOX), Erythromycin (ERM), Gentamicin (GEN), Chloramphenicol (CHL), Ciprofloxacin (CIP), Levofloxacin (LVX), and Nitrofurantoin (NIT). Briefly, Antibiotic disks were prepared by aliquoting 10 uL of antibiotic stock (GEN 1 mg/ml, AMP 1 mg/ml, CIP 0.5 mg/ml, LVX 0.5 mg/ml, ERM 1.5 mg/ml, CHL 3 mg/ml, NIT 30 mg/ml, DOX 3 mg/ml) onto the disk. Vehicle control disks were prepared similarly. Strains were streaked from glycerol stocks onto CHROMagar and incubated overnight at 37°C. Single isolated colonies were inoculated into 3 mL Brain-Heart-Infusion broth and incubated for 16 – 18 hours, then normalized to 0.5 McFarland standard, washed and resuspended in sterile 1X Phosphate-Buffered Saline (PBS). 150 µL of standardized culture were pipetted onto 150_mm BHI Agar plates and spread using sterile glass beads. Plates were dried before disks were placed on the agar. *Escherichia coli* ATCC25922 was used for quality and vehicle controls. Plates were incubated inverted overnight and antimicrobial susceptibility was evaluated by measurement of the zone of inhibition per the established zone diameter breakpoints of Clinical and Laboratory Standards Institute (CLSI) (73-75).

All Ciprofloxacin phenotypes and intermediate or resistant Gentamicin and Chloramphenicol phenotypes were further validated using Minimum Inhibitory Concentration (MIC) microdilution assay in BHI broth per CLSI breakpoints. MIC was measured using the HT-MIC workflow to determine the minimum inhibitory concentration of antibiotic required to inhibit 90% of the growth of untreated controls (MIC90) as previously described (76). In brief, 96-well 10x antibiotic master plates were prepared from antibiotic stocks using the Opentrons OT2 robot, and 30 µL was transferred to each well from the master plate to the test plates. Strains were first grown shaking at 200 rpm overnight at 37°C in a Innova S44 orbital shaker. The next day, 50 µL of each overnight culture was inoculated into fresh BHI and grown shaking at 200 rpm at 37°C until mid-log phase was reached, as determined by an OD600nm of approximately 0.4 or ~$10^8$ CFU/mL. Mid-log phase cultures were diluted to an OD600 of 0.002 (~$10^5$ CFU/mL) and 270 µL of each culture was added to a separate well of the 96-well test plate already containing 30 µL of dispensed antibiotic. The test plates were covered with lids to prevent evaporation and were incubated for approximately 20 hours shaking at 200 rpm at 37°C. After incubation, the test plates were removed, and the OD600 of each well was measured on a Biotek Synergy H1 plate reader, using BHI as a blank. Measured OD600 values were used for MIC90 determination.

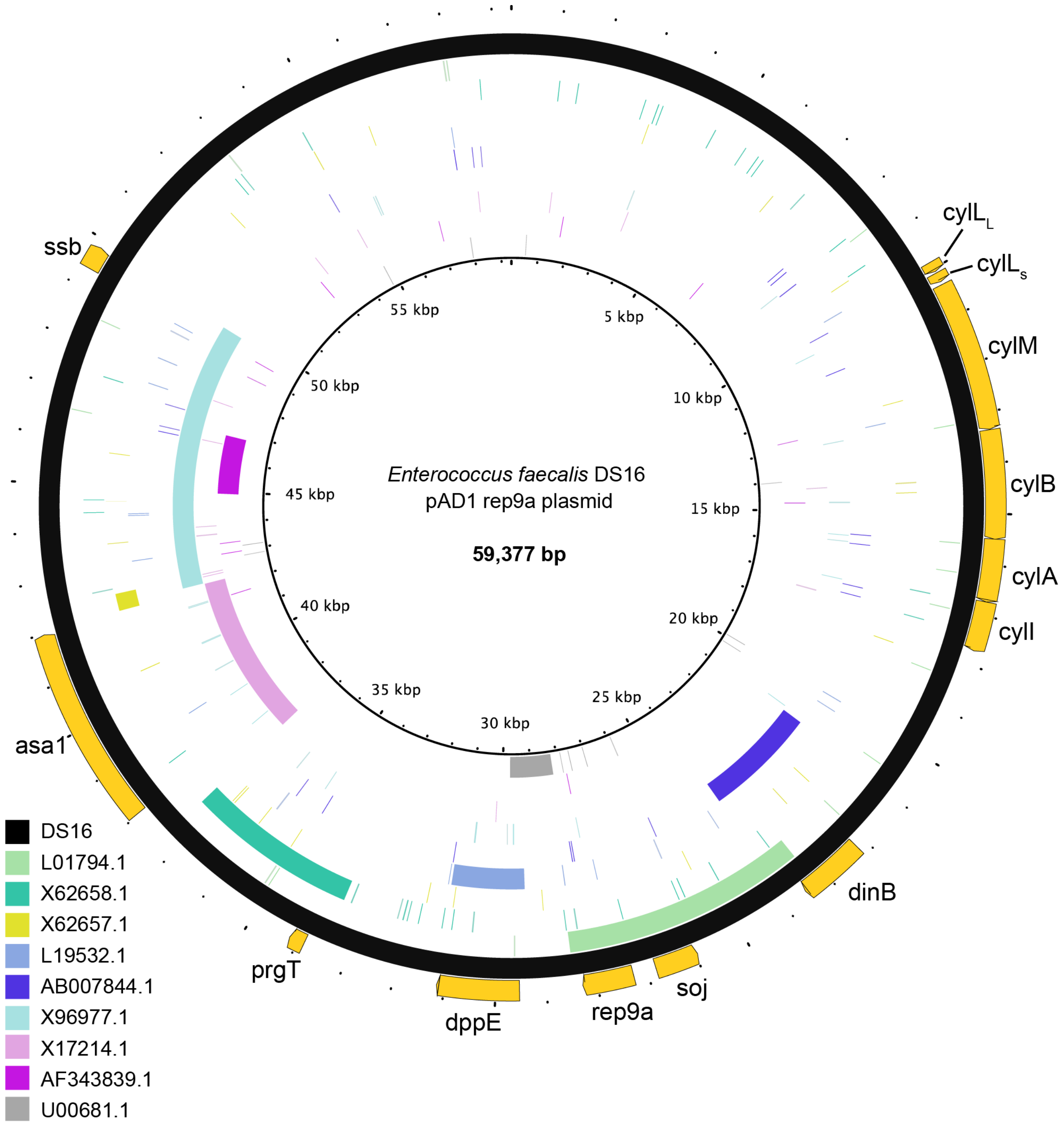


**Figure S1. Sequence alignments of complete DS16 pAD1 plasmid with previously published pAD1 sequence fragments.** BRIG blastn alignment of 9 reference pAD1 fragments (accession numbers listed in legend). Annotated genes are depicted in yellow on outermost circle. Alignment sequence identity >70%.

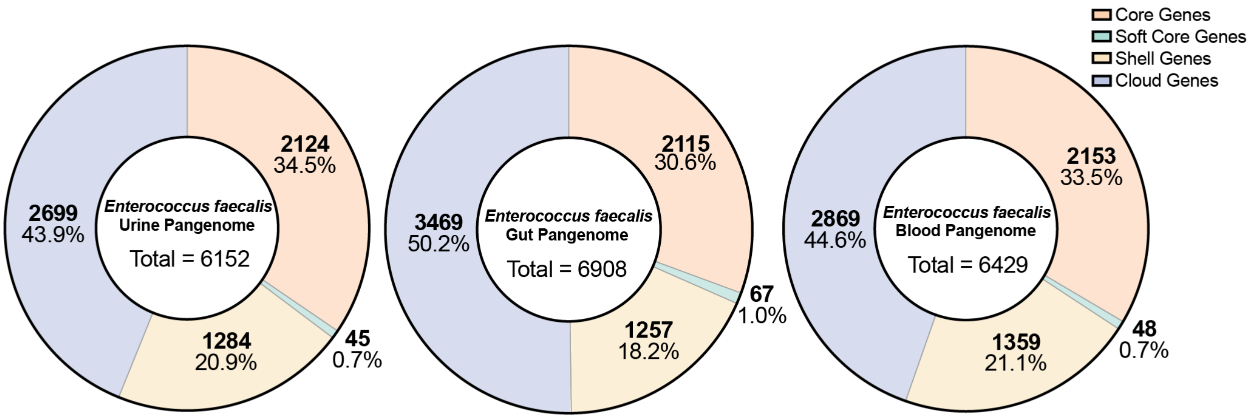


**Figure S2. Pangenomes of urinary, gut, and blood isolation groups.** Pangenome was determined for isolates in each isolation group. Core genes present in >99% of isolates, Soft Core genes present in 95-99% of isolates, Shell genes present in 15-95% of isolates, and Cloud genes present in <15% of isolates.

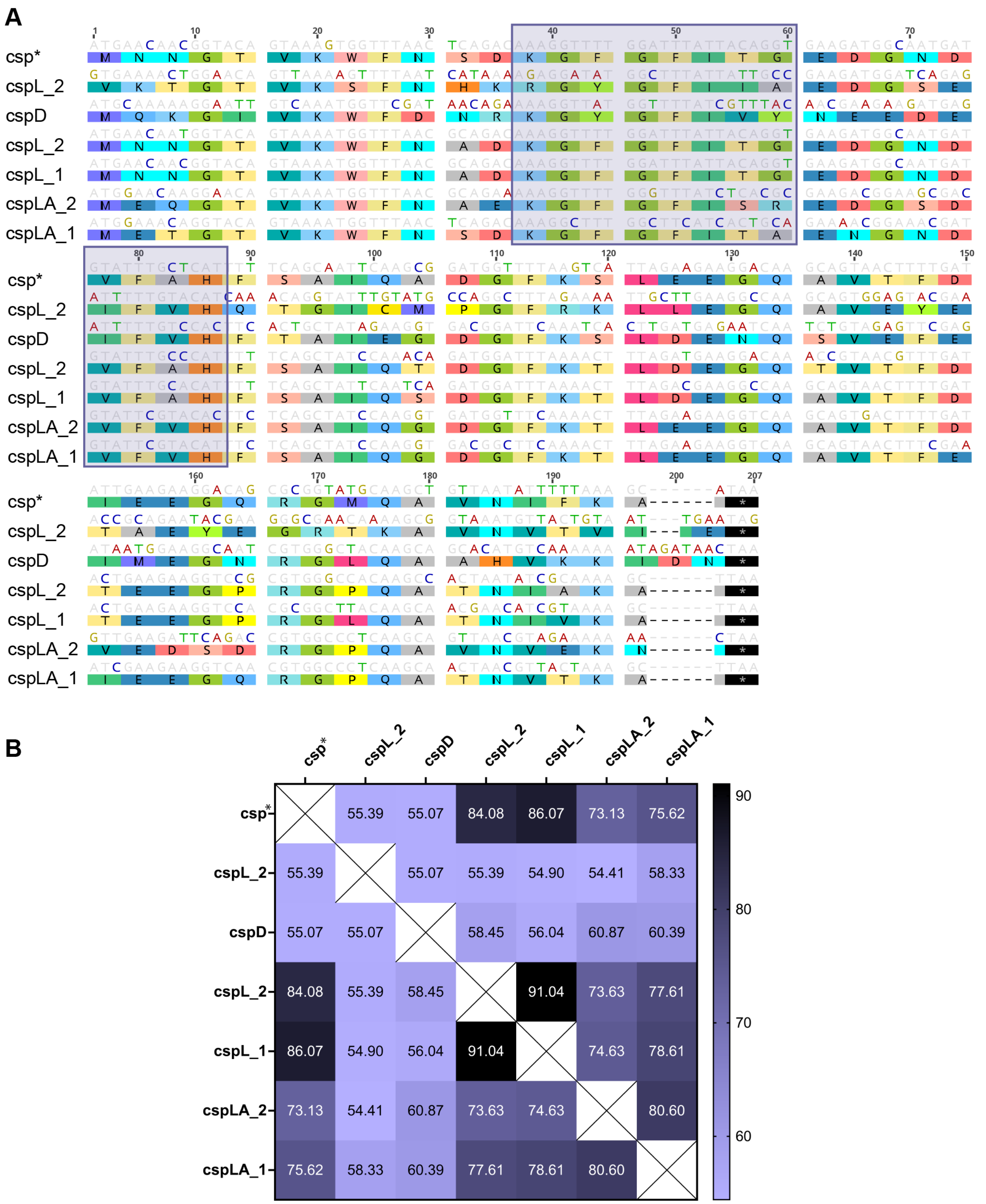


**Figure S3. Nucleotide alignment of all *csp* alleles identified in *E. faecalis*. (A)** MUSCLE nucleotide alignment of 7 *csp* alleles. Residues of nucleic acid binding (ribonucleoprotein, RNP, sites) are shaded. **(B)** Sequence identity (%) matrix of all *csp* alleles. Identity values are listed within heat map. *Enrichment candidate allele.

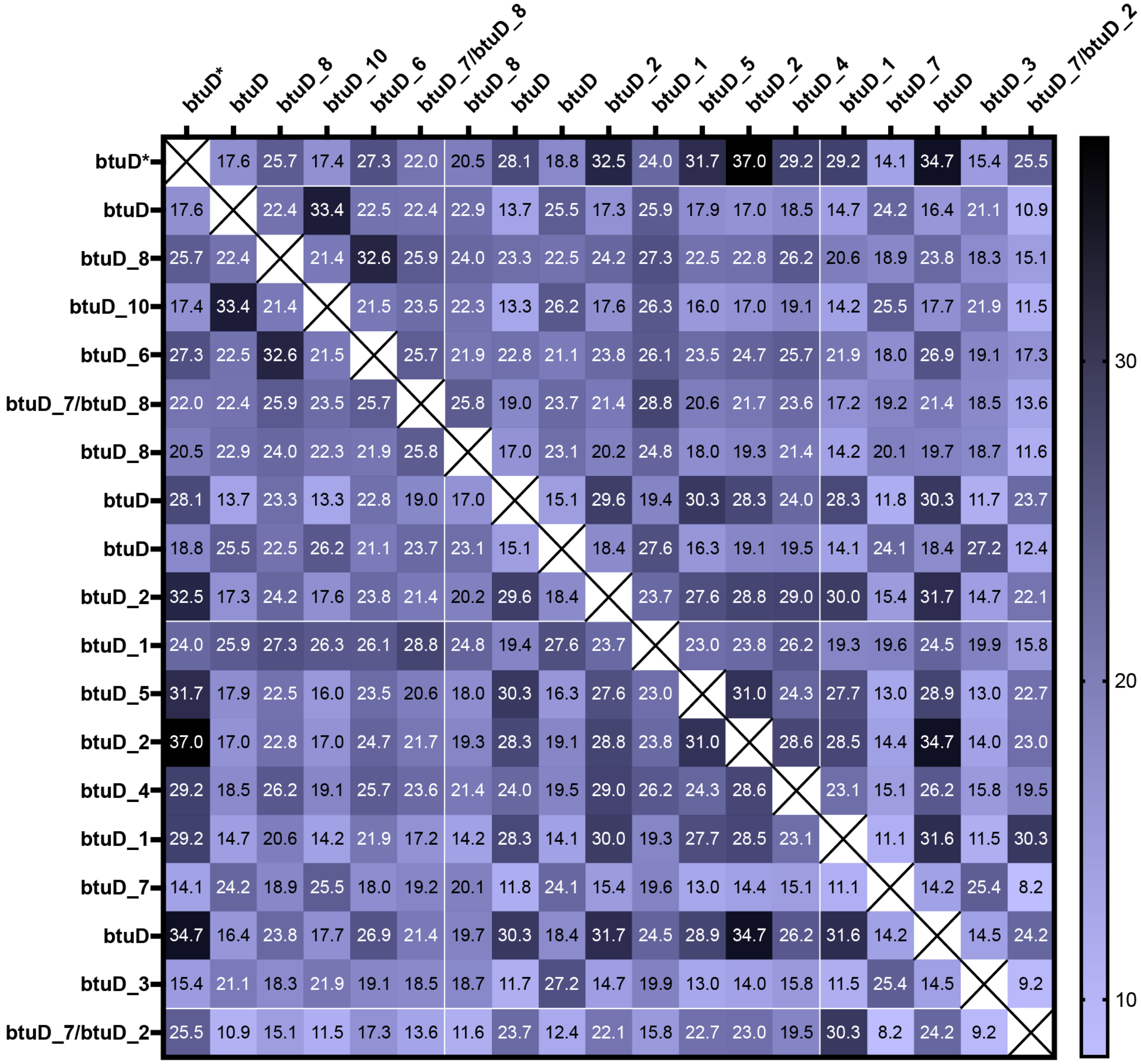


**Figure S4. Nucleotide identity (%) matrix of all *btuD* alleles identified in *E. faecalis.*** MUSCLE nucleotide alignments of representative sequences of 19 *btuD* alleles identified in the pangenome of *E. faecalis* suggest the enriched allele is a unique vitamin B12 import ATP binding protein. Identity (%) is annotated. *Enrichment candidate allele.
